# Gene model for the ortholog of *raptor* in *Drosophila eugracilis*

**DOI:** 10.64898/2026.07.10.737807

**Authors:** Anne E. Backlund, Jonas Nielsen, John Pulford, Janelle Suriaga, Jordan Pyle, Shadrian McDaniel, Jeffrey S. Thompson, Chinmay P. Rele, Jacqueline K. Wittke-Thompson

## Abstract

Gene model for the ortholog of *raptor* in the D. eugracilis Apr. 2013 (BCM-HGSC/Deug_2.0) (DeugGB2) Genome Assembly (GenBank Accession: GCA_000236325.2) of *Drosophila eugracilis*. This ortholog was characterized as part of a developing dataset to study the evolution of the Insulin/insulin-like growth factor signaling pathway (IIS) across the genus *Drosophila* using the Genomics Education Partnership gene annotation protocol for Course-based Undergraduate Research Experiences.

## Introduction

This article reports a predicted gene model generated by undergraduate work using a structured gene model annotation protocol defined by the Genomics Education Partnership (GEP; thegep.org) for Course-based Undergraduate Research Experience (CURE). The following information in quotes may be repeated in other articles submitted by participants using the same GEP CURE protocol for annotating Drosophila species orthologs of Drosophila melanogaster genes in the insulin signaling pathway.

“In this GEP CURE protocol students use web-based tools to manually annotate genes in non-model *Drosophila* species based on orthology to genes in the well-annotated model organism fruit fly *Drosophila melanogaster*. The GEP uses web-based tools to allow undergraduates to participate in course-based research by generating manual annotations of genes in non-model species (Rele et al., 2023). Computational-based gene predictions in any organism are often improved by careful manual annotation and curation, allowing for more accurate analyses of gene and genome evolution (Mudge and Harrow 2016; Tello-Ruiz et al., 2019). These models of orthologous genes across species, such as the one presented here, then provide a reliable basis for further evolutionary genomic analyses when made available to the scientific community.” (Myers et al., 2024).

“The particular gene ortholog described here was characterized as part of a developing dataset to study the evolution of the Insulin/insulin-like growth factor signaling pathway (IIS) across the genus *Drosophila*. The Insulin/insulin-like growth factor signaling pathway (IIS) is a highly conserved signaling pathway in animals and is central to mediating organismal responses to nutrients (Hietakangas and Cohen 2009; Grewal 2009).” (Myers et al., 2024).

“raptor positively regulates (Target of Rapamycin) TOR-mediated cell apoptosis and growth control by differentially regulating S6K-dependent signaling pathways, and is a crucial regulator of cell growth and metabolism in *Drosophila* (Lee and Chung 2007; Hatfield et al., 2015). *raptor* is orthologous to the human *RPTOR* gene (*regulatory associated protein of raptor complex 1*) and is a well conserved component of the TORC1 complex (Wang et al., 2012).” (Backlund et al., 2025).

“*D. eugracilis* is part of the *melanogaste*r species group within the subgenus *Sophophora* of the genus *Drosophila* (Pélandakis et al., 1993). It was first described as *Tanygastrella gracilis* by Duda (1924) and revised to *Drosophila eugracilis* by Bock and Wheeler (1972). *D. eugracilis* is found in humid tropical and subtropical forests across southeast Asia (https://www.taxodros.uzh.ch, accessed 1 Feb 2023).” (Morgan et al., 2022).

We propose a gene model for the *D. eugracilis* ortholog of the *D. melanogaster* raptor (*raptor*) gene. The genomic region of the ortholog corresponds to the uncharacterized protein XP_017064130.1 (Locus ID LOC108103241) in the D. eugracilis Apr. 2013 (BCM-HGSC/Deug_2.0) (DeugGB2) Genome Assembly of *D. eugracilis* (GCA_000236325.2 – Chen et al., 2014). This model is based on RNA-Seq data from *D. eugracilis* (PRJNA63469) and *raptor* in *D. melanogaster* using FlyBase release FB2024_02 (GCA_000001215.4; Gramates et al., 2022; Jenkins et al., 2022; Larkin et al., 2021).

### Synteny

The reference gene, *raptor*, occurs on chromosome X in *D. melanogaster* and is nested by *Multiple inositol polyphosphate phosphatase 2* (*Mipp2*), flanked upstream by *Ca2+-channel protein* _α_*1 subunit T* (*Ca-*_α_*1T*) and *Neprilysin 1* (*Nep1*), and flanked downstream by *CG4660* and *CG4666*. The *tblastn* search of *D. melanogaster* raptor-PB (query) against the *D. eugracilis* (GenBank Accession: GCA_000236325.2 Genome Assembly (database) placed the putative ortholog of *raptor* within scaffold AFPQ02004929 (AFPQ02004929.1) at locus LOC108103241 (XP_017064130.1)— with an E-value of 0.0 and a percent identity of 79.27%. Furthermore, the putative ortholog is nested by LOC108103242 (XP_017064132.1), which corresponds to *Mipp2* in *D. melanogaster* (E-value: 0.0; identity: 86.34% as determined by *blastp*; Figure 1A, Altschul et al., 1990). The putative ortholog is flanked upstream by LOC108103196 (XP_041674378.1) and LOC108103207 (XP_017064086.1), which correspond to *Ca-*_α_*1T* and *Nep1* in *D. melanogaster* (E-value: 0.0 and 0.0; identity: 89.10% and 93.95%, respectively, as determined by *blastp*). Additionally, LOC108103207 appears to be nesting LOC108103208 (XP_017064087.1), however *tblastn* did not map any *D. melanogaster* CDS to this locus, and upon a *blastp* search, no peptide in *D. melanogaster* was found to align with its protein sequence. Therefore we conclude that LOC108103208 is an ORF with misaligned RNA-Seq data and does not exist in *D. eugracilis*. The putative ortholog of *raptor* is flanked downstream by LOC108103198 (XP_017064075.1) and LOC108103202 (XP_017064080.1), which correspond to *CG4660* and *CG4666* in *D. melanogaster* (E-value: 7e-173 and 2e-144; identity: 90.42% and 99.48%, respectively, as determined by *blastp*). The putative ortholog assignment for *raptor* in *D. eugracilis* is supported by the following evidence: The genes surrounding the *raptor* ortholog are orthologous to the genes at the same locus in *D. melanogaster* and local synteny is completely conserved, supported by E-values and percent identities, so we conclude that LOC108103241 is the correct ortholog of *raptor* in *D. eugracilis* (Figure 1A).

**Figure 1.**
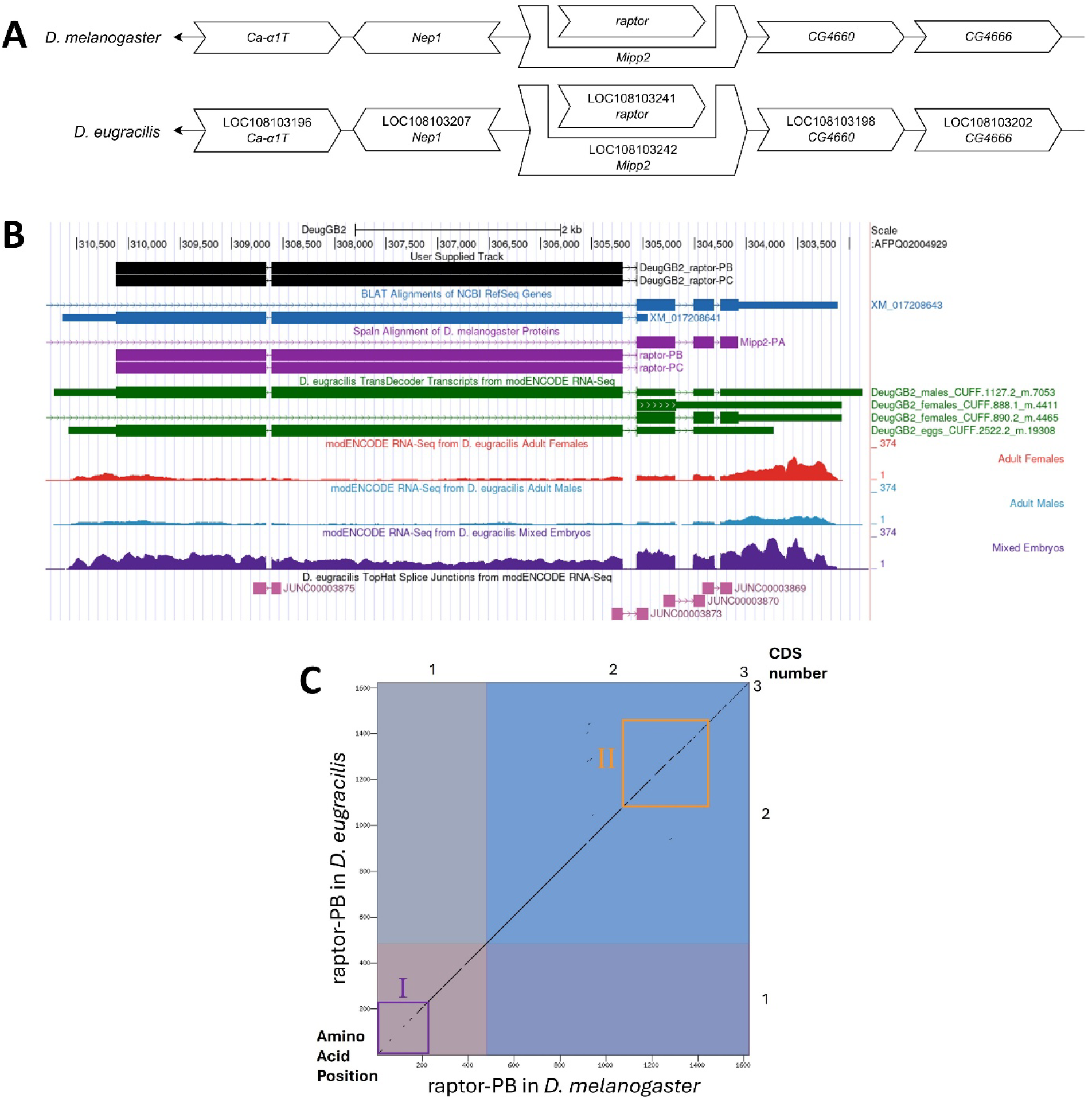
*raptor* gene model comparison between *Drosophila eugracilis* and *Drosophila melanogaster* orthologs. **(A) Synteny comparison of the genomic neighborhoods for *raptor* in *Drosophila melanogaster* and *D. eugracilis***. Thin underlying arrows indicate the DNA strand within which the target gene–*raptor*–is located in *D. melanogaster* (top) and *D. eugracilis* (bottom). The thin arrow pointing to the left indicate that *raptor* is on the negative (-) strand in *D. melanogaster* and *D. eugracilis*. The wide gene arrows pointing in the same direction as *raptor* are on the same strand relative to the thin underlying arrows, while wide gene arrows pointing in the opposite direction of *raptor* are on the opposite strand relative to the thin underlying arrows. White gene arrows in *D. eugracilis* indicate orthology to the corresponding gene in *D. melanogaster*. Gene symbols given in the *D. eugracilis* gene arrows indicate the orthologous gene in *D. melanogaster*, while the locus identifiers are specific to *D. eugracilis*. **(B) Gene Model in GEP UCSC Track Data Hub** (Raney et al., 2014). The coding-regions of *raptor* in *D. eugracilis* are displayed in the User Supplied Track (black); coding CDSs are depicted by thick rectangles and introns by thin lines with arrows indicating the direction of transcription. Subsequent evidence tracks include BLAT Alignments of NCBI RefSeq Genes (dark blue, alignment of Ref-Seq genes for *D. eugracilis*), Spaln of D. melanogaster Proteins (purple, alignment of Ref-Seq proteins from *D. melanogaster*), Transcripts and Coding Regions Predicted by TransDecoder (dark green), RNA-Seq from Adult Females, Adult Males, and Mixed Embreyos (red, light blue, and dark purple, respectively; alignment of Illumina RNA-Seq reads from *D. eugracilis*), and Splice Junctions Predicted by regtools using *D. eugracilis* RNA-Seq (PRJNA63469). The splice junctions pertaining to the *raptor* ortholog (JUNC00003875 and JUNC00003873) shown in pink have read-depths of 170 and 161, respectively. **(C) Dot Plot of raptor-PB in *D. melanogaster* (*x*-axis) vs. the orthologous peptide in *D. eugracilis* (*y*-axis)**. Amino acid number is indicated along the left and bottom; CDS number is indicated along the top and right, and CDSs are also highlighted with alternating colors. Line breaks in the dot plot indicate mismatching amino acids at the specified location between species. Two regions lack sequence similarity and are highlighted by the purple box labeled I and the orange box labeled II.

### Protein Model

*raptor* in *D. eugracilis* has three CDSs within the genome sequence. The protein sequence (raptor-PB and raptor-PC) is translated from two mRNA isoforms that differ in their UTRs (*raptor-RB* and *raptor-RC*; Figure 1B). Relative to the ortholog in *D. melanogaster*, the CDS number and protein isoform count are conserved. The sequence of raptor-PB in *D. eugracilis* has 87.29% identity (E-value: 0.0) with the protein-coding isoform raptor-PB in *D. melanogaster*, as determined by *blastp* (Figure 1C). Unusual characteristics of this model include two regions which lack sequence similarity between *D. melanogaster* and *D. eugracilis* (purple box I and orange box II; Figure 1C), and low RNA-Seq data throughout CDSs of the *raptor* ortholog. Coordinates of this curated gene model are stored by NCBI at GenBank/BankIt (accession **<u>BK064555)</u>**. This gene model can also be seen within the target genome at this <u>TrackHub</u>.

### Special characteristics of the protein model

#### Low RNA-Seq Data

There is relatively low RNA-Seq data for the *raptor* ortholog in *D. eugracilis*, with peaks occurring downstream of the 3’ end of the gene (Figure 1B). These peaks can be attributed to the untranslated region of the *Mipp2* ortholog, the nesting gene of *raptor*. In the future, long-read RNA-Seq data can be used to further investigate *raptor* in *D. eugracilis*.

## Methods

“Detailed methods including algorithms, database versions, and citations for the complete annotation process can be found in Rele et al. (2023). Briefly, students use the GEP instance of the UCSC Genome Browser v.435 (https://gander.wustl.edu; Kent et al., 2002; Navarro Gonzalez et al., 2021) to examine the genomic neighborhood of their reference IIS gene in the *D. melanogaster* genome assembly (Aug. 2014; BDGP Release 6 + ISO1 MT/dm6). Students then retrieve the protein sequence for the *D. melanogaster* reference gene for a given isoform and run it using *tblastn* against their target *Drosophila* species genome assembly on the NCBI BLAST server (https://blast.ncbi.nlm.nih.gov/Blast.cgi; Altschul et al., 1990) to identify potential orthologs. To validate the potential ortholog, students compare the local genomic neighborhood of their potential ortholog with the genomic neighborhood of their reference gene in *D. melanogaster*. This local synteny analysis includes at minimum the two upstream and downstream genes relative to their putative ortholog. They also explore other sets of genomic evidence using multiple alignment tracks in the Genome Browser, including BLAT alignments of RefSeq Genes, Spaln alignment of *D. melanogaster* proteins, multiple gene prediction tracks (e.g., GeMoMa, Geneid, Augustus), and modENCODE RNA-Seq from the target species. Detailed explanation of how these lines of genomic evidenced are leveraged by students in gene model development are described in Rele et al. (2023). Genomic structure information (e.g., CDSs, intron-exon number and boundaries, number of isoforms) for the *D. melanogaster* reference gene is retrieved through the Gene Record Finder (https://gander.wustl.edu/~wilson/dmelgenerecord/index.html; Rele et al., 2023). Approximate splice sites within the target gene are determined using *tblastn* using the CDSs from the *D. melanogaste*r reference gene. Coordinates of CDSs are then refined by examining aligned modENCODE RNA-Seq data, and by applying paradigms of molecular biology such as identifying canonical splice site sequences and ensuring the maintenance of an open reading frame across hypothesized splice sites. Students then confirm the biological validity of their target gene model using the Gene Model Checker (https://gander.wustl.edu/~wilson/dmelgenerecord/index.html; Rele et al., 2023), which compares the structure and translated sequence from their hypothesized target gene model against the *D. melanogaster* reference gene model. At least two independent models for a gene are generated by students under mentorship of their faculty course instructors. Those models are then reconciled by a third independent researcher mentored by the project leaders to produce the final model. Note: comparison of 5’ and 3’ UTR sequence information is not included in this GEP CURE protocol.” (Gruys et al., 2025)

## Supporting information

Gene model data files

## Supplemental Files

1. Zip file containing a FASTA, PEP, GFF files for the gene model
2. Figure 1 in high resolution

### Metadata

Bioinformatics, Genomics, *Drosophila*, Genotype Data, New Finding

## Acknowledgements

We would like to thank Wilson Leung for developing and maintaining the technological infrastructure that was used to create this gene model and Laura K. Reed for overseeing the project. Thank you to <u>FlyBase</u> for providing the definitive database for *Drosophila melanogaster* gene models. Further, we would like to thank the editors and developers at the journal *microPublication: Biology* for assistance in developing the template for these single gene ortholog publications.

